# Enhanced Deep Convolutional Neural Network for SARS-CoV-2 Variants Classification

**DOI:** 10.1101/2023.08.09.552643

**Authors:** Mike J. Mwanga, Hesborn O. Obura, Mudibo Evans, Olaitan I. Awe

**Author notes:** Corresponding author: Olaitan I. Awe.

## Abstract

High-throughput sequencing techniques and sequence analysis have enabled the taxonomic classification of pathogens present in clinical samples. Sequencing provides an unbiased identification and systematic classification of pathogens and this is generally achieved by comparing novel sequences to pre-existing annotated reference databases. However, this approach is limited by large-scale reference databases which require considerable computational resources and skills to compare against. Alternative robust methods such as machine learning are currently employed in genome sequence analysis and classification, and it can be applied in classifying SARS-CoV-2 variants, whose continued evolution has resulted in the emergence of multiple variants.

We developed a deep learning Convolutional Neural Networks-Long Short Term Memory (CNN-LSTM) model to classify dominant SARS-CoV-2 variants (omicron, delta, beta, gamma and alpha) based on gene sequences from the surface glycoprotein (spike gene). We trained and validated the model using > 26,000 SARS-CoV-2 sequences from the GISAID database. The model was evaluated using unseen 3,057 SARS-CoV-2 sequences. The model was compared to existing molecular epidemiology tool, nextclade.

Our model achieved an accuracy of 98.55% on training, 99.19% on the validation and 98.41% on the test dataset. Comparing the proposed model to nextclade, the model achieved significant accuracy in classifying SARS-CoV-2 variants from unseen data. Nextclade identified the presence of recombinant strains in the evaluation data, a mechanism that the proposed model did not detect.

This study provides an alternative approach to pre-existing methods employed in the classification of SARS-CoV-2 variants. Timely classification will enable effective monitoring and tracking of SARS-CoV-2 variants and inform public health policies in the control and management of the COVID-19 pandemic.

## 1. Introduction

Taxonomic classification of pathogens in clinical samples is an important step in the diagnosis and tracking of outbreaks. Conventional methods such as polymerase chain reactions (PCR) and metagenomic analyses have been applied in the diagnosis and taxonomic classification of organisms isolated from clinical samples. PCR approaches employ primers designed to specifically bind to targeted genomic regions of pathogens [1,2]. This approach requires prior information on the genomic structure of the pathogens for the design and manufacture of primers. The design of primers and probes limits these methods to identify only known variants and require continuous updates to cater for newer species or strains that are clinically relevant. High-throughput sequencing provides an unbiased identification of microbes such as bacteria, viruses, parasites and fungi molecules present in clinical samples and identification of novel microbes [3] and identification of disorders in newborns [4]. Gene expression in plants (Die *et al*., 2019) and pathogen evolution using SARS-CoV-2 [5] have also been studied using Next-generation sequencing data.. These methods are capable of identifying and characterizing pathogens without prior knowledge of a specific pathogen from clinical samples [6]. Next-generation sequencing and analysis has gained strides in the recent past due to its reduced cost, improved sequencing speed, more rapid and availability of user-friendly analysis tools allowing its application in the wider microbial research [7]. Commonly, this technique evaluates sequence similarities between novel query sequences and predetermined reference sequences [8,9]. Classifying a large number of short sequence-reads generated by sequencing technologies takes much time and is computationally intensive [10]. This approach is also hindered by the lack of well-curated and annotated metagenomic reference sequences to classify newly discovered species [11]. Additionally, the exponential increase in the number of sequenced microbes has increased the level of comparisons for newly identified microbes resulting in large-scaled reference databases of microbes to compare against, and considerable computational challenges [10]. Furthermore, bioinformatics analysis of the partial or whole genome sequences requires highly skilled expertise; limiting timely identification of the circulating pathogens. Consequently, this results in prolonged clinical turnaround time for prognosis leading to delays in monitoring outbreaks.

Recent advancements in technology such as machine learning (ML), and other artificial intelligence techniques have revolutionized medicine and genomics [12]. Machine learning based tools can identify and extract important sequence features for sequence classification in a computationally efficient manner. In a DNA sequence, these features will be the pattern of arrangement of nucleotides in a sequence that is/are unique for each pathogen [8]. For this reason, machine learning methods have been applied in the identification and classification of pathogens in clinical samples. For instance, VirFinder [13] models used *k-mer* frequency to classify viruses while VirSorter [14] uses a probabilistic model tool to predict viral sequences. Randhawa *et al*., (2020) proposed a supervised machine learning with a digital signal process (MLDSP) model for genomic identification of CoVID-19 virus signatures, important in the classification of SARS-CoV-2 [15]. Despite machine learning models showing strides in the classification of pathogens, they are still faced with a major drawback; their inability to automatically extract important hidden features from DNA sequences and have to rely on manual feature extraction which can be laborious and computationally intensive.

The recent emergence of deep learning, a subclass of machine learning, has offered solutions to these challenges. Deep learning approach uses natural language processing to represent data at high abstraction levels based on multiple non-linear transforming layers with excellent results in automated feature extraction [8,16]. Deep learning has also been applied extensively in DNA classification, gene, and protein structural predictions, and other genomic analysis projects including viral identification and classification. Within the infectious disease discipline, deep learning algorithms such as VirHunter [17], DeepVirFinder [18] and ViralMiner [19] which rely on a combination of convolutional neural networks and dense neural networks have been applied in the identification of viral sequences from raw metagenomic datasets. Additionally, PACIFIC algorithm has been developed to identify respiratory viral co-infections occurring in CoVID-19 patients from RNA-seq data. Deepthi *et al*., 2021 has utilized a deep learning ensemble approach to screen clinically approved antiviral drugs for potential efficacy against the novel SARS-CoV-2 and the eventual drug repurposing against COVID-19 disease. Such and many more deep learning methods address challenges related to homology-dependent methods and swiftly detect and classify pathogenic sequences [20].

The most common deep-learning model that can produce cutting-edge results for most classification problems is Convolutional Neural Networks (CNN) [16,21]. CNN models are known to produce high accuracy in image and text data categorization such as spam detection and sentiment classification [16]. As such researchers have made use of this model to study images from medical and public health studies. For instance, [22] and [23] (Balaram *et al*., 2022) proposed a CNN model that detects and classifies malaria parasites from blood smears. CNN models have also been applied in determining the presence of CoVID-19 before reaching a mass-scale level in patients using CT-scan images [24]. Additionally, chest X-ray images from patients with CoVID-19 have been used to categorize disease severity classes; mild, moderate, severe and critical [25]. According to work by [26] CNNs have been applied in agriculture for the classification of various plant diseases using infected plant leaf images, which achieved an accuracy of 91%. In a study by Tharsanee *et al*., (2021) [24], a deep learning approach was applied to classify mango leaves infected with fungal diseases known as anthracnose, the model achieved an accuracy of 96.16%. In most machine learning models, users are required to select features for training the model. In contrast, CNN models automatically extract features from the dataset [16]. by mapping a specific length of input to a fixed-size output and training with a back-propagation algorithm [21]. This makes the CNN architecture efficient for large datasets including genome sequences from clinical samples [15,27] and offer alternative approaches to current existing methods.

Since the beginning of CoVID-19, genetic lineages of severe acute respiratory syndrome coronavirus-2 (SARS-CoV-2) have been emerging and circulating globally. Currently, the SARS-CoV-2 Interagency Group (SIG) has categorized five SARS-CoV-2 variants of concern (VOC) namely; alpha, beta, gamma, delta, and omicron [28]. These variants were first detected in different parts of the world: alpha variant emerged in the UK, beta in S. Africa, gamma in Brazil, delta in the USA, and omicron from Botswana [29]. The emergency, spread, and transmission of SARS-CoV-2 variants is highly associated with mutations on the spike gene whose genetic characteristics are used to identify and distinguish the variants [30,31]. The surface glycoprotein gene i.e spike protein, plays a critical role in the invasion of the host by the virus; thus, the gene has been a target for many studies related to possible therapeutic targets [30,31]. Increased mutations in the spike protein results in the emergence of novel variants with increased transmission rates and overwhelming the available therapeutic measures [28,32]. Computational sequencing is used to compare genetic differences between the observed viruses and characterize variants and their relationships [33]. For instance, genomic surveillance of SARS-CoV-2 has been applied in the characterization of variants and understanding of local and global circulation and transmission patterns [34–37]. Given the continuous emergence and spread of SARS-CoV-2 variants, there is a need to develop further efficient and advanced methods for variant identification and classification to aid in the surveillance and monitoring of such and similar infectious pathogens. Here, we employ a (CNN) component of deep learning techniques, on publicly available genomics datasets to classify SARS-CoV-2 variants. This will further improve timely identification and classification to help prevent outbreaks and assist in the design and implementation of control and containment measures.

## 2. Methods

There are 5 key steps in this process; 1) data pre-processing which involves one-hot encoding and annotation, 2) feature extraction model identifies a unique pattern in the sequence, 3) convolving the features into feature maps, 4) flattening, and 5) classification.

### 2.1. Dataset

Development of the CNN model made use of publicly available representative SARS-CoV-2 glycoprotein spike gene nucleotide sequences downloaded from the GISAID database (https://www.gisaid.org/). The conventional variant names were used to download variant-specific sequences with the following filters within the GISAID databases; complete, high coverage, low coverage exclude, collection data complete, and collection dates between 01/01/2021 to 8/4/2022. Sequences were selected with the representation of pre-defined geographical regions (Africa, Asia, Oceania, S. America, N. America). Spike gene sequences were extracted using the blastn algorithm [38]. The downloaded sequences were used to create the blast database while the spike gene sequence from the Wuhan reference genome (NC_045512.2/Wuhan-Hu-1) was used as the query sequence [39]. Sequence alignment was achieved using MAFFT v7.4.5 [40]. The sequence alignment step was meant to rearrange the data into dimensions with the same data shape for both training and testing dataset. This allows the model to learn the patterns from the training data and be able to process the test data in the same manner. Figure 1 shows the distribution of gene sequence for each of the SARS-CoV-2 variants.

**Figure 1.**
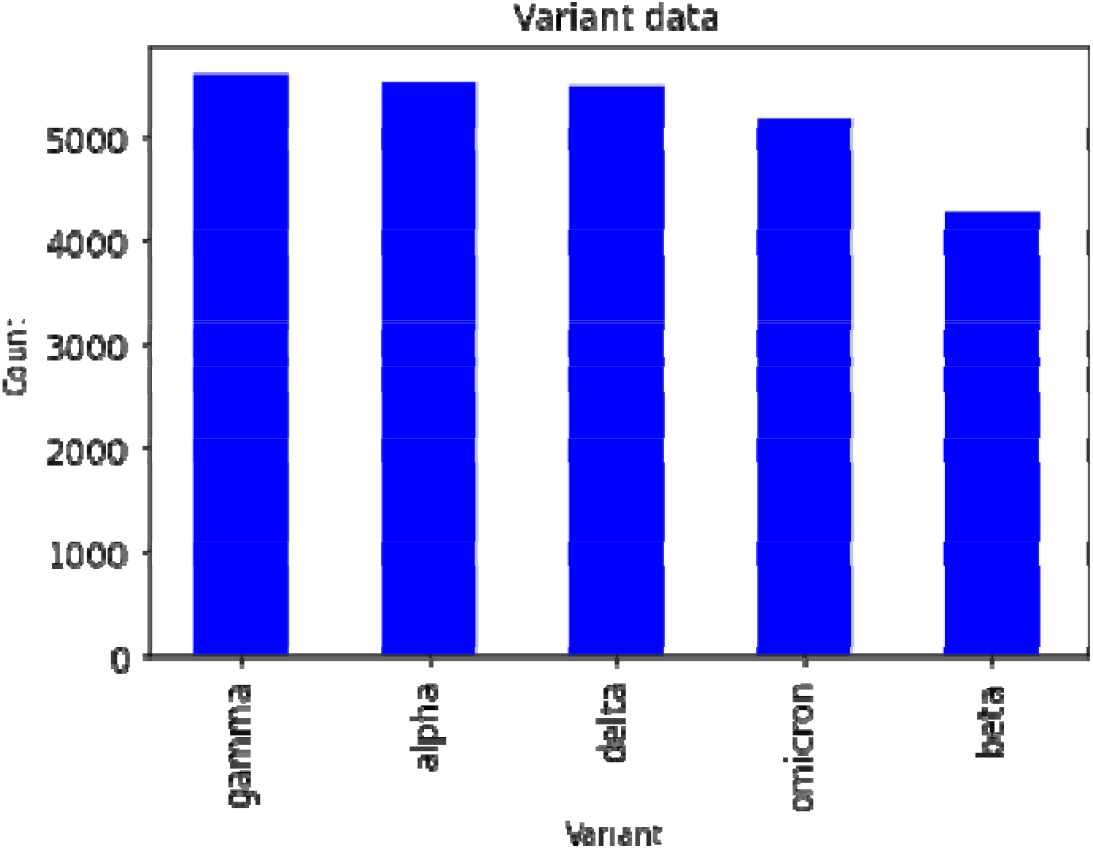
Distribution of each SARS-COV-2 variant for deep learning model

Machine learning classification task consists of five main stages; data preprocessing, feature extraction, model training, and finally classification [8,16].

### 2.2. Data preprocessing

Python libraries were used in data processing and exploratory analysis before feeding to the model. The input feature was the DNA sequence, while the output features were the names of the SARS-CoV-2 variants: alpha, beta, gamma, omicron, and delta. Machine learning algorithms make use of numerical data and since DNA sequences are in text form, nucleotide base conversion using readily available tools was implemented. One-hot encoding was manually employed to transform the DNA sequences into a machine-readable binary matrix. The four DNA bases; A, C, G, and T, were the informative characters of the dataset and hence assigned unique vectors containing ones and zeros, see figure 3. All other non-DNA characters in the sequence including ambiguous bases and gaps were considered non-informative hence assigned vectors with zeros, see figure 3. Insertions were treated as normal DNA nucleotides and assigned a binary label as indicated in the figure 3. In contrast, deletions were treated as uninformative sites similar to ambiguous bases, hence not considered during feature selection by the model. The resulting dataset was split into training (75%) and validation (25%) and fed into the CNN model, see figure 2.

**Figure 2.**
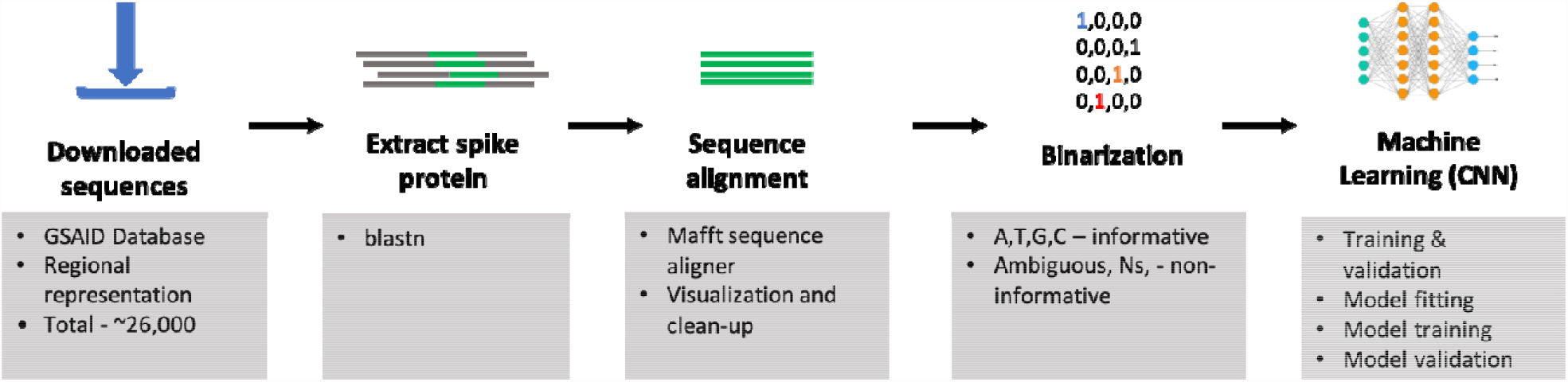
Data preprocessing workflow: feature extraction, feature encoding, and classifications. The first step is to input raw sequences for feature extraction, and feature encoding using the one-hot encoding technique. Then the one-hot encoded sequences are fed to the CNN model for classification analysis.

**Figure 3.**
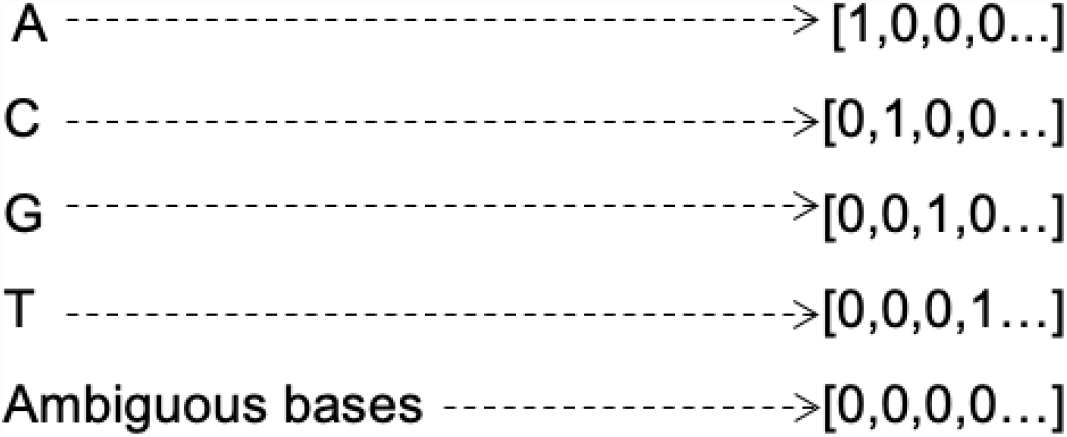
Overview of the one-hot encoding method. Binary arrays are assigned to each DNA base and non-DNA bases.

### 2.3. CNN-Long Short-term Memory (CNN-LSTM) Model Architecture

In this study, the model was developed from scratch with four convolutional layers combined with a rectified linear activation function (ReLU). The last convolution layers are connected to a bidirectional recurrent neural network [41]. The fully connected layer has 6 dense layers connected to the softmax algorithm for classification as shown in table 1.

**Table 1.** CNN architecture for SARS-COV-2 variants classification.

| Step | Operation | Output Dimension |
| --- | --- | --- |
| Input layer | One-Hot encoding | 3849x13 |
| Convolutional Layer 1 | Conv1D (54,6) | 3849x54 |
|  | Activation (ReLU) | 769x54 |
|  | Max-Pooling1D | 769x54 |
|  | Dropout (0.5) | 769x54 |
| Convolutional Layer 2 | Conv1D (27,3) | 769x27 |
|  | Activation (ReLU) | 769x27 |
|  | Max-Pooling1D | 256x27 |
| Convolutional Layer 3 | Conv1D (14,2) | 256x14 |
|  | Activation (ReLU) | 256x14 |
|  | Max-Pooling1D | 85x14 |
|  | Dropout (0.2) | 85x14 |
| Convolutional Layer 4 | Conv1D(7,2) | 85x7 |
|  | Activation (ReLU) | 85x7 |
|  | Max-Pooling1D | 28x7 |
|  | Bidirectional (512) | 1024 |
|  | Dropout (0.01) | 1024 |
| Flatten layer | Flatten | 1024 |
| Dense layer 1 | Dense (256) | 256 |
|  | Activation (ReLU) | 256 |
| Dense layer 2 | Dense (128) | 128 |
|  | Activation (ReLU) | 128 |
| Dense layer 3 | Dense (64) | 64 |
|  | Activation (ReLU) | 64 |
| Dense layer 4 | Dense (32) | 32 |
|  | Activation (ReLU) | 32 |
| Dense layer 5 | Dense (16) | 16 |
|  | Activation (ReLU) | 16 |
| Dense layer 6 | Dense (5) | 5 |
|  | Activation (SoftMax) |  |
| **Parameters** |  |  |
| Total parameters | 1,916,397 |  |
| Trainable parameters | 1,916,397 |  |

To switch from one convolutional layer to the next, CNN stores and updates its weights after learning the relationship between the input and output. Firstly, the model was initialized with random parameters during forwarding propagation and the parameters are fine-tuned during backward-propagation, minimizing the mean squared error (loss). We performed hyperparemeter tuning as the number of epochs increased, by adding more layers or removing some layers and while observing the model accuracy on model training. In this study, we employed categorical cross-entropy as a loss function.

The convolutional layer is a key step in CNN model as it does feature extraction from the one-hot encoded input layer creating a feature map that contains a particular pattern from the DNA sequences [42]. Padding is set as ‘same’ to preserve the original size of the input layer [43]. Applying a convolutional layer over the input layer results in the reduction of the output layer leading to the loss of important features. To navigate this, we pad the input layer with zeros around the border for the input vector [44]. Each convolutional layer is followed by a max-pooling layer which reduces the dimensional and complexity of the extracted features in the network. Commonly used pooling layers are maximum pooling, average pooling, stochastic pooling, and spectral [45]. In this study, the parameters and convolutional layers were empirically tested and selected. Maximum pooling was used as follows; filter size: 6×6, 3×3, 2×2, and 2×2. A dropout layer is used as a regularization technique thus preventing overfitting [46]. In this network, each convolutional layer is followed by a rectified linear unit activation function (ReLU), which introduces non-linearity that solves the vanishing gradient effect. ReLU converts every negative number from the pooling layer to zero [47]. In the flatten layer, 2-dimensional feature maps produced in the previous layer are converted into a 1-dimensional feature map to be suitable for the following fully connected layers [22]. The last layer of the convolutional layer has an LSTM layer which overlooks insignificant parts of the preceding layers and carefully updates the important feature as output that is required. This approach solves the vanishing gradient in the architecture [48,49]. The fully connected layer has several dense layers with the last layer having a softmax activation algorithm; a multiclass classifier for calculating the probabilities to which the five variants belong [22]. Adaptive moment estimation optimizer (Adam) is used as an optimizer in the neural network architecture. Adam computes the averages and squares for a gradient for each parameter [50].

This study used 30 epochs and a batch size of 1000. After every epoch, the filter weights are updated to minimize the loss function. CNN-LSTM training occurs through a back propagation technique with the ultimate goal to optimize the weights of the nodes by calculating the gradient [49]. The following steps describe the training process: random weight initialization at the input layer, forward propagation of the weights in the network where each node uses its inputs and associated weights to calculate the activation values, calculation of the loss function at the output layer, and back propagation.

This model was implemented in Python version 3.9.5 using Keras version 2.8.0; a framework for the development of neural networks.

## 3. Results

The dataset used in this study consists of 26,036 spike-protein DNA sequences. The training set contained 19,527 (75%) sequences, while the validation set contained 6,509 (25%) sequences. The model had a total of 1,916,397 trainable parameters. In an Intel Xeon Linux system with 18 cores and 512 GB RAM, it took approximately 30 minutes to train the model.

### Model testing and validation

To train the model, the matrix containing the input layer is fed into the neural network, concurrently validating the model for making predictions of SARS-CoV-2 variants. The outcome and error are figured, if the outcome is wrong then it will reinitialize the weights by backpropagation process where it propagates from output to the first convolutional layer. In this model, the weights are optimized by the Adam optimizer. The proposed model architecture can detect and classify five variants of SARs-CoV-2 with a high level of accuracy and precision. To get the desired classification accuracy 30 epochs were performed. As the gradual increase in epochs, the training and validation loss reduced significantly. The training loss was at 0.07 and the validation loss was at 0.05. The training accuracy initially was 0.21 and with an increase in the number of epochs, at the thirtieth epoch, the accuracy was 98.55% Similarly, the validation accuracy was 0.21 at the first epoch, and at the thirtieth epoch, the validation accuracy increased to 99.19% as shown in figure 4 below.

**Figure 4.**
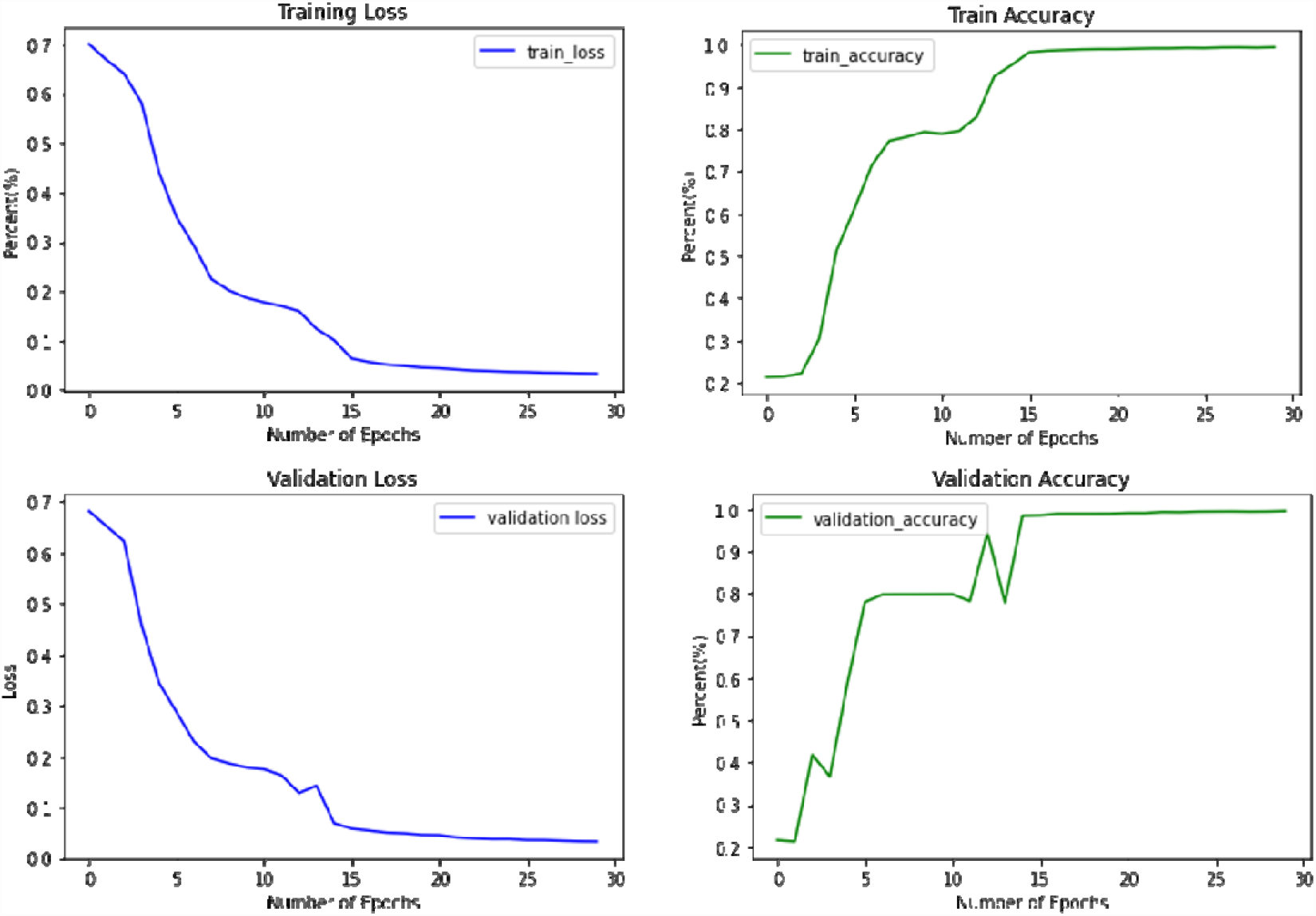
Accuracy metrics of the model on the training and validation dataset.

### 3.1. Model evaluation on new dataset

Following successful training and testing of the model, further evaluation of the model was performed on a new dataset containing 3,057 SARS-CoV-2 variant sequences downloaded from the GSAID database. The distribution of the dataset for each variant is shown in figure 5.

**Figure 5.**
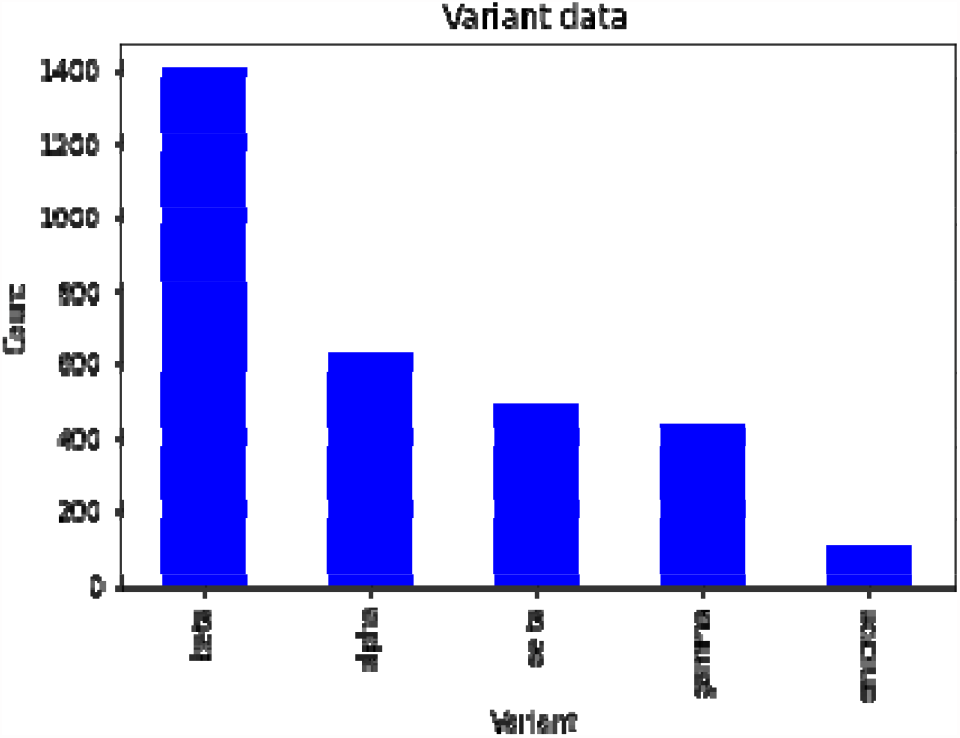
Distribution of sequences for the five SARS-CoV-2 variants in the evaluation dataset.

### 3.2. Model Evaluation

The performance of the model was evaluated from the confusion matrix shown below in figure 6. The figure depicts the overall confusion matrix of the five different SARS-CoV-2 variants from the model. It clearly shows that 432 (14%) gamma, 1396 (46%) beta, 463 (15%) delta, 103 (3%) omicron, and 616 (21%) alpha variants were correctly classified by the model. In contrast, few variants were misclassified; 9 sequences of the alpha variant were classified as beta, 2 and 2 sequences of the beta variant were classified as gamma and alpha respectively. Additionally, 28 and 3 sequences of the delta variant were classified as beta and alpha variants while 5 gamma sequences were classified as beta. This study included a confusion report which has the precision, recall, accuracy and F1 score for the 5 classes, shown in table 2 below. Precision is the ability of the model to predict the true positive sequence as positive and recall shows the number of positive sequences that are classified correctly as positive. The F1 score (F-score or F-measure) sumps up the predictive performance of the model by combining precision and recall metrics. It gives the overall accuracy of the model. This is calculated from the precision and recall test as shown in the following formulas.

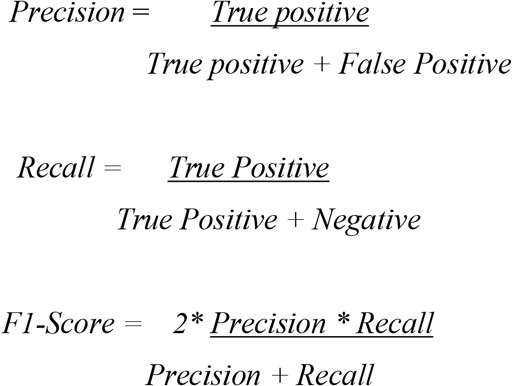

**Table 2:** Performance report of the model on unseen data.

| Metrics | Variant | Precision | Recall | F1-Score | Number of Sequences |
| --- | --- | --- | --- | --- | --- |
|  | Gamma | 1.0 | 0.99 | 0.99 | 437 |
|  | Delta | 1.0 | 0.94 | 0.97 | 494 |
|  | Beta | 0.94 | 1.0 | 0.97 | 1403 |
|  | Alpha | 1.0 | 0.98 | 0.99 | 625 |
|  | Omicron | 0.7 | 0.42 | 0.52 | 103 |
| Accuracy | - | - | - | 0.98 | 3026 |
| Weighted average | - | 0.98 | 0.98 | 0.98 | 3026 |

**Figure 6.**
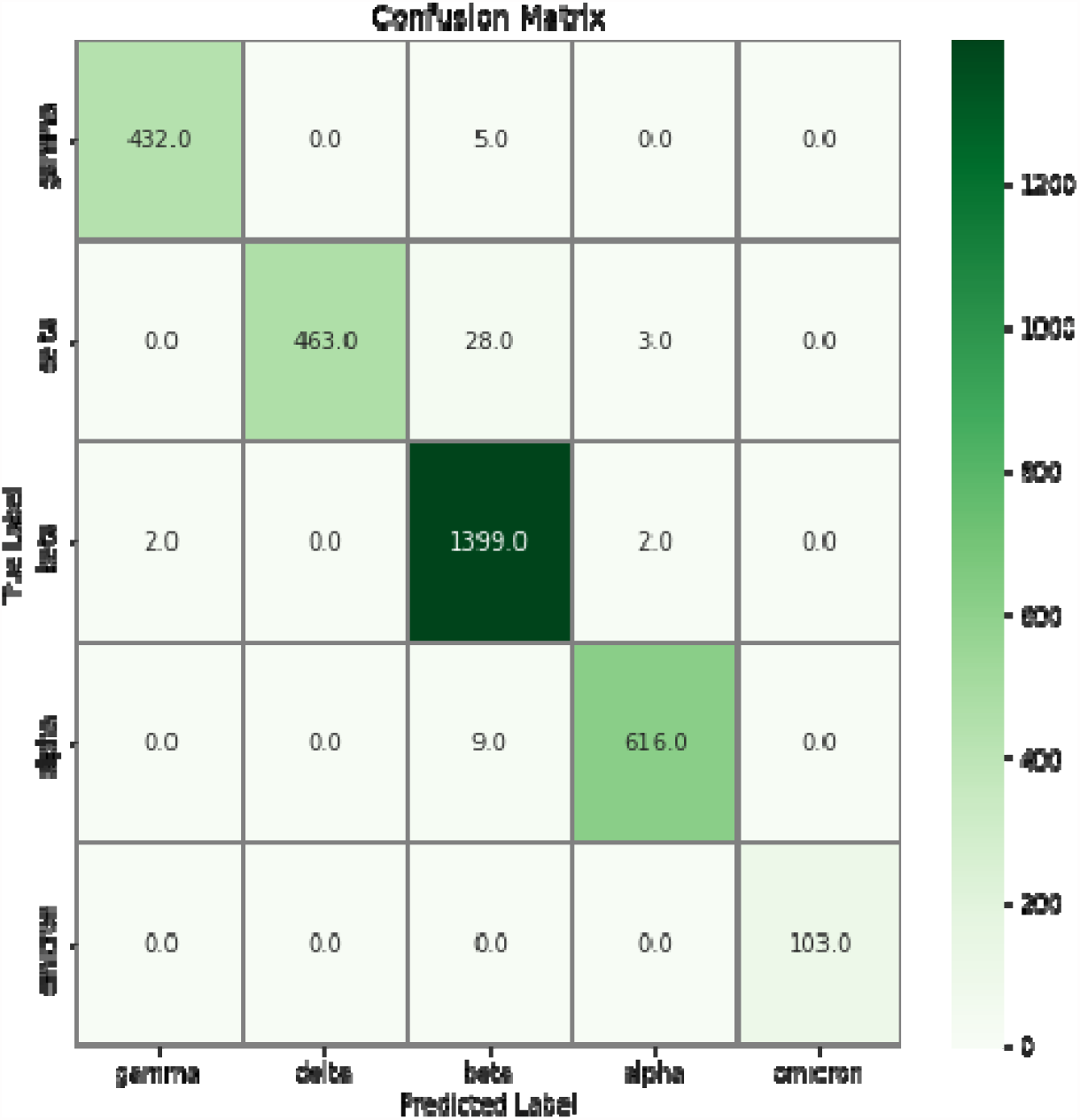
Confusion matrix showing performance of the model on new data

From the evaluation matrix, the model shows an accuracy of 98.41% given a new dataset for classification of the SARS-CoV-2 variants.

Alpha, gamma and delta variants from the confusion report have a precision of 100.00% indicating that the model obtains excellent results in the validation datasets.

We compared validation results from the model and classification results using Nextclade - a genomic analysis tool that performs, among other things, variant assignment of SARS-CoV-2 sequences [51]. Table 3 below shows the metrics of comparison between the two tools. The model showed better performance, with over 90% accuracy in variant assignment for alpha, beta, omicron and gamma variants, with a decreased accuracy (87%) for delta variant. Besides the commonly identified variants, nextclade also identified the presence of recombinant SARS-CoV-2 strains in omicron (n=67) and delta (n=11).

**Table 3:** Comparison between proposed model and nextclade in variant classification.

|  |  | Nextclade |  |  |  | CNN+LSTM Model |  |
| --- | --- | --- | --- | --- | --- | --- | --- |
| Variant | Total | Assigned | % | Unassigned | Recombinant | Assigned | Model Success % |
| Alpha | 625 | 620 | 99 | 5 | 0 | 616 | 98 |
| Beta | 1403 | 1398 | 99 | 5 | 0 | 1399 | 99 |
| Gamma | 437 | 436 | 99 | 1 | 0 | 432 | 91 |
| Delta | 494 | 482 | 98 | 1 | 11 | 463 | 87 |
| Omicron | 103 | 36 | 35 | 0 | 67 | 103 | 100 |

## 4. Discussion

In this study, we trained and evaluated a CNN-LSTM deep learning model to classify SARS-CoV-2 variants, based on the spike gene sequences.

This study demonstrated that the model is robust enough when applied to a new dataset to determine the class of a SARS-CoV-2 variant. This model achieved an accuracy of 98% in assigning unknown sequences to their respective classes. In real-world scenarios, generated reads and genomes contain multiple ambiguous bases and Ns due to sequencing errors. Such errors are challenging during alignments and complicate sequence analysis workflows. In this study, the model architecture disregards the presence of these ambiguous bases and any Ns present in the gene sequences as zeros (not informative). As such, the model only utilizes and extracts features based on present DNA bases, making it an excellent and reliable tool for real-world sequencing analysis studies. Secondly, the high accuracy achieved by this model can be attributed to the use of data from the spike gene, which is only ∼13% of the complete genome. Comparing the proposed model to pre-existing molecular epidemiology tools such as nextclade, our model achieves significant accuracy in classifying SARS-CoV-2 variants from unseen data. However, nextclade identified the presence of recombinant strains in the evaluation data, a mechanism that the proposed model did not detect. We aim to subsequently reinforce this model by incorporating more features to distinguish diversity mechanisms such as recombination events, lineages and novel variants, for improved classification. This approach reduces the number of unnecessary features encompassed within the genome and computes complexity in modeling whole genome data hence higher accuracy in classification. Furthermore, since deep learning models are data intensive and require much time to train, the use of over 20,000 gene sequences to train the model and the availability of computational resources for more train-time increases the urgent need to deploy machine learning models to complement clinical diagnosis and variants classification. Integrating such models in clinical setup helps to reduce clinical turn-around time for SARS-CoV-2 variant determination. This will help inform public health policies on containment measures.

To our knowledge, this is the first CNN architecture designed to classify SARS-CoV-2 variants using features from spike gene sequences. Other machine learning models have been employed in genomic studies in different paradigms, achieving excellent results. For instance, (Whata & Chimedza, 2021) proposes a CNN algorithm including a bi-directional long short term memory (Bi-LSTM) for the classification of SARS-CoV-2 viruses within the Coronavirus family. This model achieved an accuracy of 99.95% and an area under curve receiver operating characteristic (AUC ROC) of 100.00%. However, the lack of sufficient computational resources meant that this model was not developed on deeper architectures and large datasets, limiting confidence in the proposed architecture. In another study by [52] authors present KEVOLVE, a machine learning model based on a genetic algorithm to determine SARS-CoV-2 variant discriminative signatures using complete genome sequences. With a precision of 0.99, KEVOLVE outperformed other multiple statistical tools in identifying key motifs used to differentiate SARS-CoV-2 variants. Such studies are encouraging enough to recognize the significance of deep learning as an alternative to analysis of genomics (omics) sequence data and support surveillance programs.

## 5. Conclusion

This study shows that deep learning can be applied as an alternative method to the classification of viruses in addition to conventional sequence classification methods. CNN-LSTM model achieved high accuracy in classifying the five most dominant SARS-CoV-2 variants. This tool could be deployed in the context of the current CoVID-19 pandemic in assisting researchers with real-time surveillance and monitoring of circulating coronavirus strains.

## Acknowledgements

The authors thank the National Institutes of Health (NIH) Office of Data Science Strategy (ODSS) and the National Center for Biotechnology Information (NCBI) for their immense support before and during the April 2022 Omics codeathon organized in collaboration with the African Society for Bioinformatics and Computational Biology (ASBCB). The authors acknowledge Daniel Lumian for helping in editing the first draft of the manuscript.

## Author Information

## Contributions

MJM and HOO conceived the original idea. MJM and HOO developed the ml_sarscov2 pipeline, performed the bioinformatic analysis of the case study data, and drafted the manuscript. MJM and HOO intensively tested the pipeline and provided feedback. EM assisted in writing the original draft of the manuscript. OIA provided supervision, reviewed the manuscript and provided critical feedback and resources that were needed in order to successfully do the project and required analysis. All authors read and approved the final manuscript.

## Ethics approval and consent to participate

Not applicable.

## Data Availability

Data and code used in this study are available in this GitHub repository (https://github.com/omicscodeathon/ml_sarscov2).

## Consent for publication

Not applicable.

